# The protein architecture of the endocytic coat analyzed by FRET

**DOI:** 10.1101/480996

**Authors:** Michal Skruzny, Emma Pohl, Sandina Gnoth, Gabriele Malengo, Victor Sourjik

**Affiliations:** Department of Systems and Synthetic Microbiology and 35043 Marburg, Germany; Flow Cytometry and Imaging Facility, Max Planck Institute for Terrestrial Microbiology, 35043 Marburg, Germany; LOEWE Center for Synthetic Microbiology (SYNMIKRO), 35043 Marburg, Germany.

**Keywords:** endocytosis, clathrin, membrane reshaping, FRET, yeast

## Abstract

Endocytosis is a fundamental cellular trafficking pathway, which requires an organized assembly of the multiprotein endocytic coat to pull the plasma membrane into the cell. Although the protein composition of the endocytic coat is known, its functional architecture is not well understood. Here we determine the nanoscale organization of the endocytic coat by FRET microscopy in yeast. We assessed proximities of 18 conserved coat-associated proteins and used clathrin subunits and protein truncations as molecular rulers to obtain a high-resolution protein map of the coat. Furthermore, we followed rearrangements of coat proteins during membrane invagination and their binding dynamics at the endocytic site. We show that the endocytic coat is stratified into several functional layers situated above and below the clathrin lattice with key proteins transversing through the lattice deeply into the cytoplasm. We propose that this conserved design enables an efficient and regulated function of the endocytic coat during endocytosis.

## Introduction

The knowledge of the nanoscale organization of cellular machines is crucial for understanding their function and mechanism of action. This certainly applies to the endocytic machinery, which is responsible for nutrient uptake, signaling and homeostasis in all eukaryotic cells (McMahon and Boucrot, 2011; Mettlen et al., 2018). The endocytic machinery consists of dozens of proteins, which sequentially assemble beneath the plasma membrane, at the endocytic site, to aid in endocytic site initiation, cargo selection, plasma membrane invagination, and finally vesicle scission and uncoating (Kaksonen and Roux, 2018).

The functionally paramount protein assembly of the endocytic site is the endocytic coat complex made of the clathrin lattice and multiple endocytic adaptor and scaffold proteins. The endocytic coat is essential for setting up the endocytic site, for cargo selection, and for many membrane-reshaping steps of endocytosis (Kirchhausen et al., 2014; Merrifield and Kaksonen, 2014). Initially, the clathrin lattice was thought to be the main structural element of the coat, possibly implying its organization on adaptor proteins, which connect it to the plasma membrane. The observation of rapid rearrangements of the clathrin lattice at the endocytic site has challenged this view (Wu et al., 2001; Avinoam et al., 2015; Scott et al., 2018). Moreover, other studies indicated that endocytic adaptors form regular assemblies on the plasma membrane independently of clathrin (Skruzny et al., 2015; Garcia-Alai et al., 2018). These examples illustrate that we still have very limited information about the protein organization of the endocytic coat. This is, however, imperative for understanding the function of the endocytic coat during endocytosis.

The analysis of the protein architecture of the endocytic coat is a challenging task due to its complex, compact and dynamic character. Even the rather simple endocytic coat of yeast *Saccharomyces cerevisiae* contains more than 20 proteins (each in 10-90 copies) densely packed in a hemisphere of 30-50 nm radius and associated together for a limited time of 20-40 seconds (Boettner et al., 2012; Kukulski et al., 2012; Weinberg and Drubin, 2012; Goode et al., 2015). Hence, while recent correlative electron and super-resolution light microscopy studies brought progress in the understanding the lateral organization of the endocytic site (Sochacki et al., 2017; Mund et al., 2018), description of the functional architecture of the endocytic coat is still missing.

Importantly, densities, copy numbers, and ratios of coat proteins at the endocytic site seem to be suited for mapping their organization by Förster resonance energy transfer (FRET). FRET specifically occurs between fluorophores separated by less than 10 nm and therefore allows the detection of nanometer distances between fluorescently-tagged molecules *in vivo*. FRET has already been successfully used to map the organization of several protein machines (e.g. kinetochore, spindle pole body, contractile ring, chemotactic sensory system) (Miller et al., 2005; Kentner and Sourjik, 2009; Aravamudhan et al., 2014; Gryaznova et al., 2015; McDonald et al., 2017).

Here, we apply FRET to determine the nanoscale architecture of the multiprotein endocytic coat. We analyzed 227 protein-protein proximities of all 18 conserved coat-associated proteins at the yeast endocytic site by FRET, and followed their binding dynamics and rearrangements by FRAP and FRET, respectively. We found that endocytic coat proteins localize not only inside the clathrin lattice, but organize themselves in functional layers on both sites of the clathrin meshwork. Moreover, several critical endocytic proteins transverse the lattice to connect the membrane and cytoplasmic areas of the coat. This design has extensive mechanistic and regulatory implications for the function of conserved endocytic coat during endocytosis.

## Results

### Comprehensive FRET-based protein proximity screen of the yeast endocytic coat

The yeast endocytic coat is built within the diffraction-limited endocytic site by the sequential assembly of 10-90 copies of **~** 20 coat proteins. Although the *in vivo* organization of such a supramolecular assembly cannot be explored by standard fluorescence microscopy, it can be efficiently attained by FRET-based protein proximity mapping. This approach is based on the fact that FRET occurs only between fluorescently-tagged proteins separated by less than 10 nm, and its presence thus clearly reports their proximity. To analyze proximities between individual coat-associated proteins we used the acceptor photobleaching (donor de-quenching) FRET technique. Here, a specific increase of fluorescence of the FRET donor attached to the first protein, happening after photobleaching of the FRET acceptor on the second protein, signalizes the presence of FRET and proximity between the proteins. Contrary to FRET methods based on the sensitized acceptor emission, this technique is very robust and straightforward, as it does not require extensive corrections and directly provides FRET efficiency (FRET%) values. We focused our proximity mapping on the late stage of endocytic coat formation when all known coat-associated proteins are present (Weinberg and Drubin, 2012). For that, we stabilized late endocytic coats on the flat plasma membrane by addition of Latrunculin A (LatA). LatA allows the full assembly of the endocytic coat, but blocks the subsequent actin polymerization-dependent membrane invagination steps of endocytosis (Kaksonen et al., 2003; Newpher et al., 2005; Kukulski et al., 2012).

We first constructed yeast strains expressing tandems of coat-associated proteins tagged with GFP and mCherry from their endogenous loci. We used GFP and mCherry as FRET donor and acceptor, respectively, as they were repeatedly shown not to disturb coat proteins expression and function (Kaksonen et al., 2003, 2005; Boeke et al., 2015) and to be a good FRET pair at the same time (Albertazzi et al., 2009; Aravamudhan et al., 2014). We C-terminally tagged all 14 endocytic coat proteins conserved between yeast and human (human homologs in parentheses): Syp1 (FCHo1/2); Ede1 (Eps15/R); Apl1 subunit of AP-2 complex (AP-2); Yap1801, Yap1802 (CALM/AP180); Ent1, Ent2 (epsins 1-3); Sla2 (Hip1R); Pan1, Sla1 (intersectin-s); End3 (Eps15/R); clathrin subunits Chc1 and Clc1 (CHC, CLC) and Gts1 (small ArfGAP). We also tagged 4 conserved coat-associated regulators of actin polymerization at the endocytic site: Las17 (WASP/N-WASP), Vrp1 (WIP), Bzz1 (syndapin or FCHSD1-2) and Lsb3 (SH3YL1). Additionally, we tagged selected coat-associated proteins on their N-termini with GFP in strains containing partner protein with the C-terminal mCherry tag. The only combinations not assessed were protein pairs tagged both N-terminally due to a non-functional N-terminal mCherry tag on our proteins (data not shown). Altogether, we constructed 217 GFP-mCherry protein pair strains covering 91% of the 237 biologically functional and technically accessible combinations (see Supplementary Figure S1).

To estimate the maximal FRET values achievable in our system we first analyzed FRET between GFP-mCherry tandem C-terminally appended to Sla1 and Sla2 proteins (Khmelinskii et al., 2012). We measured mean FRET efficiency of 15.1 ± 1.3% and 18.8 ± 1.4% (FRET% with 95% confidence intervals), respectively, which were indeed the highest values of all our FRET measurements. We then analyzed FRET at endocytic sites of LatA-treated cells expressing individual GFP/mCherry-tagged protein pairs. We obtained 44 FRET-positive protein pairs ranging in FRET efficiencies between 1.1-8.9%. No FRET was detected for other 173 protein pairs and for all donor-only expressing controls. The obtained FRET data are summarized in Figure 1 and detailed in Supplemetary Figure 1. The comparison of GFP donor fluorescence before and after mCherry photobleaching for 5 selected protein pairs is shown in Figure 2 and Movie 1.

**Figure 1.**
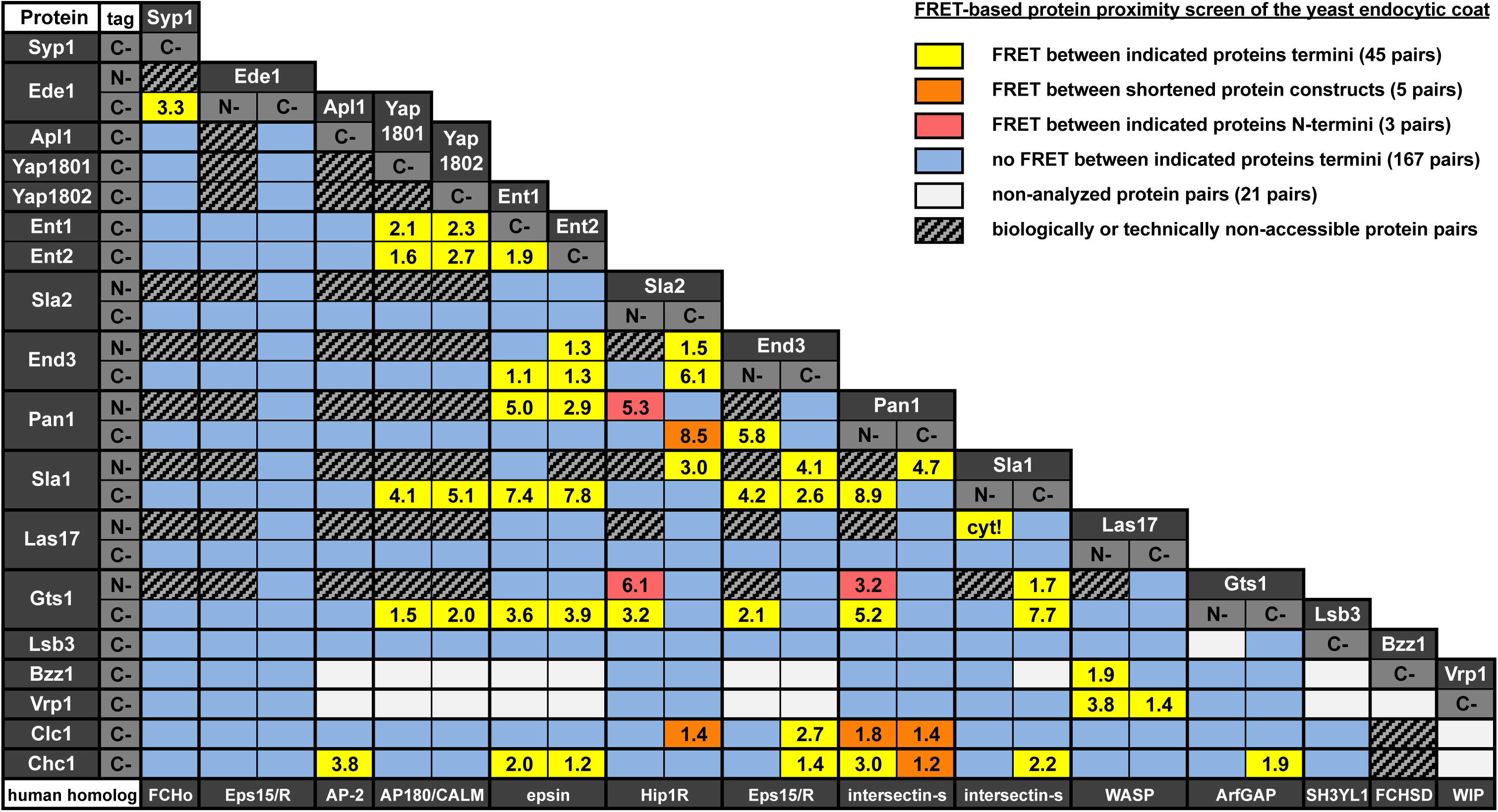
Results of FRET-based proximity screen of the yeast endocytic coat. Mean FRET efficiencies (in %) are shown for FRET-positive protein pairs (yellow, orange, and red boxes). “cyt!” indicates FRET between Las17 and Sla1 specifically detected in the cytoplasm. See legend on the top right for further details. For detailed results see Supplementary Figure 1.

**Figure 2.**
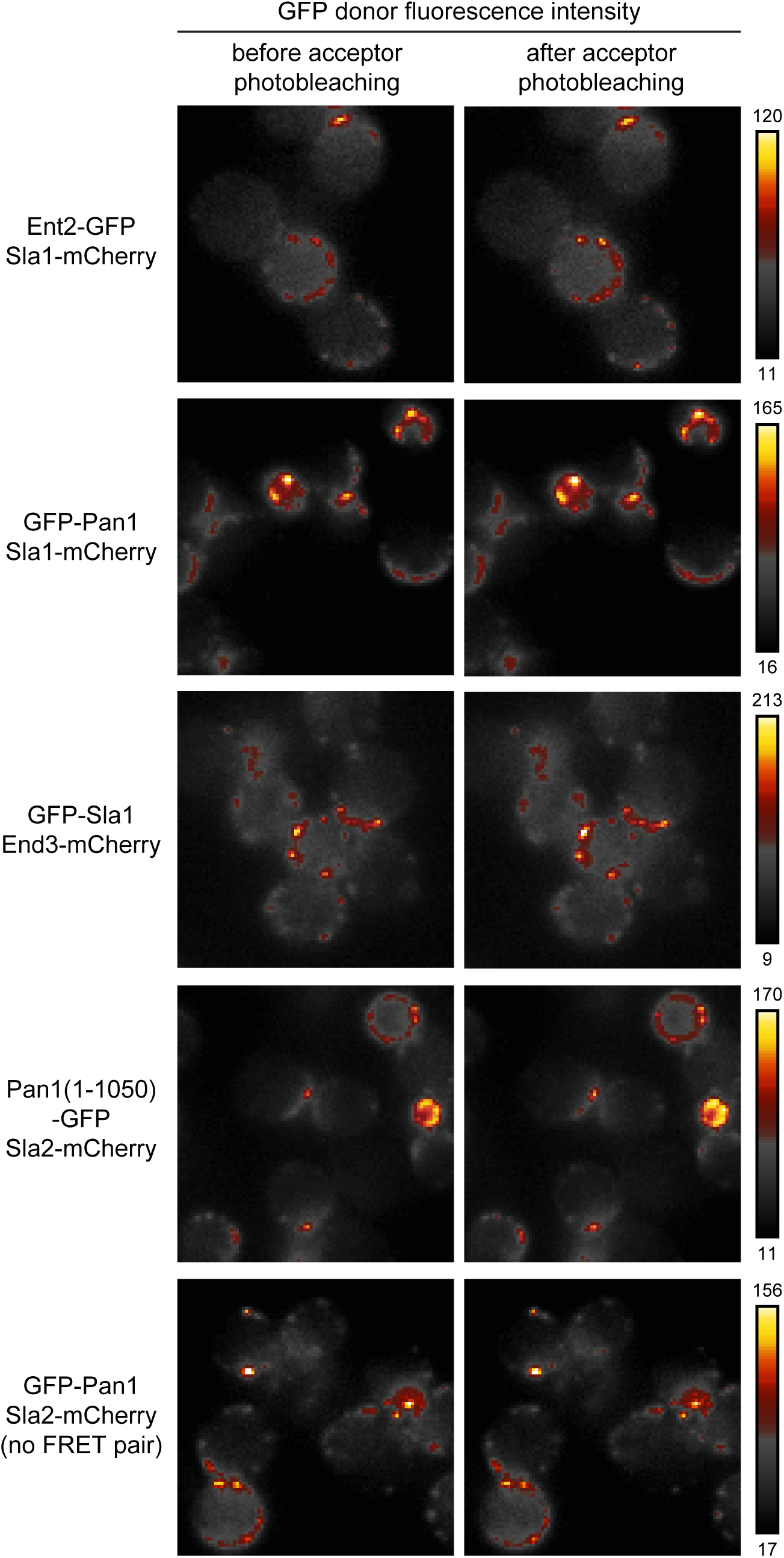
Examples of FRET proximities between endocytic coat proteins. Fluorescence intensity of the FRET donor (GFP) is shown before and after photobleaching of the FRET acceptor (mCherry) in ImageJ Smart pseudo-color scheme (for intensity values see bars on the right). FRET is observed as an increase of donor fluorescence after acceptor photobleaching.

### Building the protein map of the endocytic coat

#### Endocytic coat contains two separate protein proximity networks

We next aimed to draw the protein map of the yeast endocytic coat based on the FRET proximities detected in our screen (Figure 3A). We oriented our map towards the plasma membrane, and used two minimal constrains for the initial protein placement: first, it is thought that N-terminal membrane-binding domains of Syp1, Apl1, Yap1801/2, Ent1/2, and Sla2 are adjacent to the membrane. The N-terminus of Sla2, which can be functionally tagged with GFP (Picco et al., 2015) and was shown to be ∼1.5 nm offset of the membrane *in vitro* (Skruzny et al., 2015), can therefore serve as a topological marker for the plasma membrane proximity. Second, similarly to its human homolog Hip1R, Sla2 likely forms long dimers (Yang et al., 1999; Engquist-Goldstein et al., 2001) with their C-termini remote of the plasma membrane. In agreement, previous live-cell imaging of Sla2 tagged on both ends showed Sla2 as a **~** 33 nm-long protein, with its N-terminus close to the membrane and its C-terminus extended to the cytoplasm (Picco et al., 2015).

**Figure 3.**
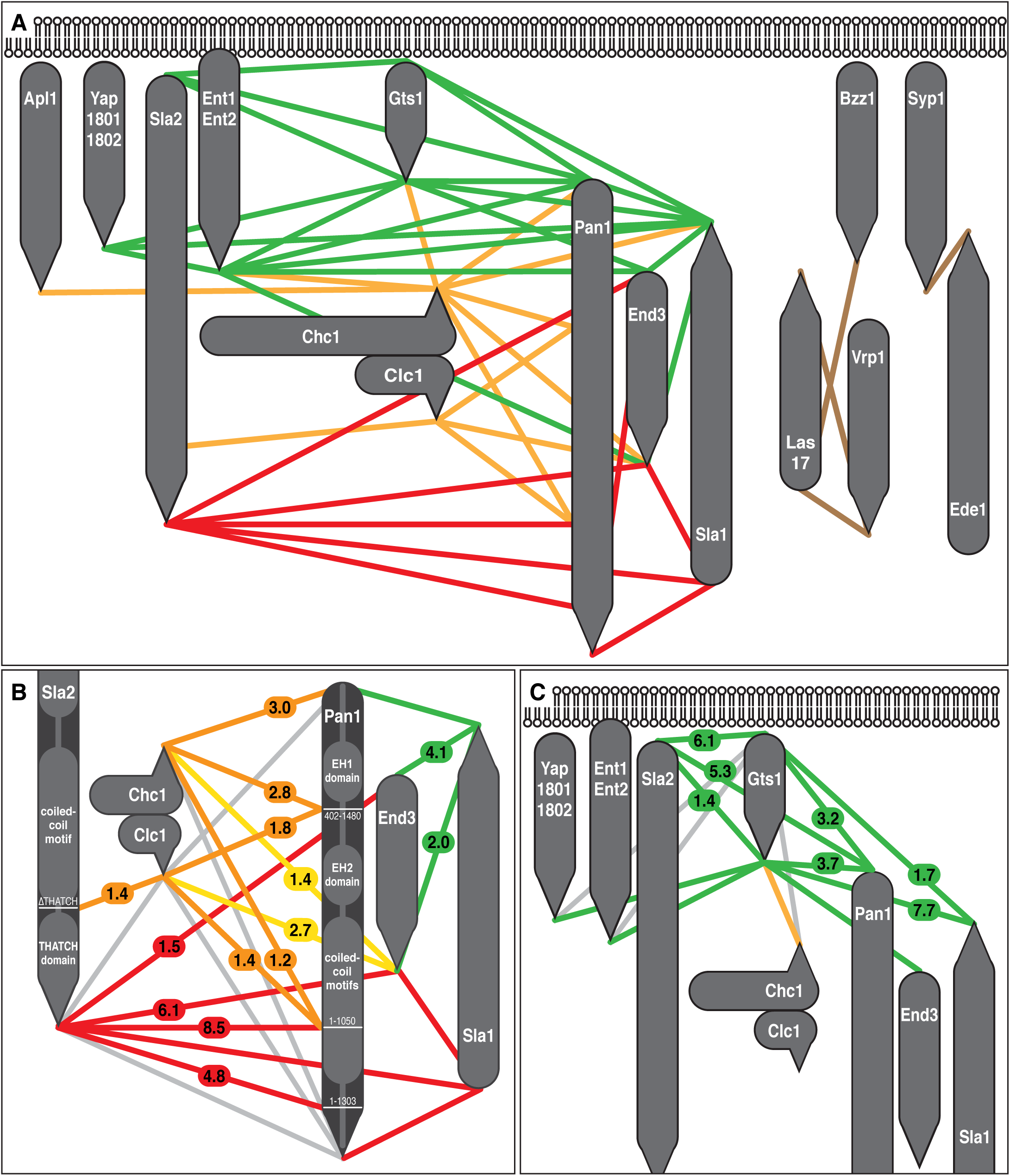
Protein map of the yeast endocytic coat based on detected FRET proximities of coat-associated proteins. **(A)** Protein proximity map of the endocytic coat-associated proteins. **(B)** Enlarged view of the map focused on selected proximities of clathrin subunits, End3, and Sla2 and Pan1 shortened and full-length proteins. Enlarged view of the map focused on Gts1 proximity network. Rounded and pointed ends of gray rods represent the N-and C-termini of indicated proteins, respectively. Single rods are used for Yap1801/2 and Ent1/2 protein duplicates, which have almost identical set of proximity partners (see Supplementary Figure S1). Membrane-proximal, clathrin-associated and cytoplasmic proximity networks are colored green, orange, and red, respectively. Isolated protein proximities are shown in brown. Gray lines in **(B)** and **(C)** show selected protein pairs showing no FRET. Yellow lines in **(B)** highlight proximities of the End3 C-terminus to the clathrin subunits. Numbers report mean FRET efficiencies.

When we connected the termini of protein pairs showing FRET, we observed three isolated FRET pairs and two remarkably cross-linked proximity networks made of several proteins termini (green and red networks in Figure 3A). The first proximity network was made of Yap1801/2, Ent1/2, Sla1, Gts1, and End3 C-termini and N-termini of Pan1 and End3. The second proximity network consisted of N-termini of Sla1 and End3, and C-termini of Sla2, Pan1, and End3. As these two proximity networks exclusively contained the opposite ends of long modular proteins Sla2, Pan1 and Sla1, we speculated that their constituents occupy distinct areas of the endocytic coat.

#### Endocytic coat proteins localize on both sites of the clathrin lattice

Next we focused on FRET proximities of all proteins to the C-terminally tagged subunits of clathrin, Chc1 and Clc1 (orange lines in Figure 3A). According to the known structure of the clathrin lattice, the C-termini of Chc1 and Clc1 are exposed on opposite surfaces of the lattice, serving thus as topological markers for areas between the lattice and the plasma membrane (Chc1) and between the lattice and the cytoplasm (Clc1). We detected the Apl1, Ent1/2, End3, Sla1, and Gts1 C-termini, and the Pan1 N-terminus as the specific FRET partners of Chc1. This strongly suggests that constituents of the first proximity network are situated between the plasma membrane and the clathrin lattice.

The only proximity to Clc1 was found for the End3 C-terminus with its FRET value twice higher than for Chc1 (2.7% vs. 1.4%; yellow lines in Figure 3B). Interestingly, two other proteins showed FRET with both End3 termini: the Sla1 C-terminus, a member of Chc1-proximal network, showed higher FRET with the End3 N-terminus than with the End3 C-terminus (4.1% vs. 2.0%, *p*=0.022, *n*=11 cells; Figure 3B). Inversely, the Sla2 C-terminus showed lower FRET with the End3 N-terminus than with the End3 C-terminus (1.5% and 6.1%; *p*<0.001, *n*=8 cells; Figure 3B). Altogether, this was suggestive of overall proximity of End3 to the clathrin lattice with its N-terminus closer to the first proximity network, and its C-terminus closer to Clc1 and the second network made of the Sla2 and Pan1 C-termini and the Sla1 N-terminus. These termini could therefore localize at the cytoplasmic site of the clathrin lattice, supposedly too remote to show FRET with Clc1.

We tested this hypothesis by measuring FRET between clathrin subunits and C-terminally shortened Pan1 and Sla2 proteins (The Sla1 N-terminus cannot be shortened without a functional loss.). We deleted C-terminal, 430 amino-acid long, unstructured part of Pan1 (Pan1(1-1050)); and C-terminal, supposedly **~** 7 nm long, THATCH domain of Sla2 (Brett et al., 2006). Both deletions were previously shown to retain essential functions, proper localization and expression levels of respective wild-type proteins (Wesp et al., 1997; Skruzny et al., 2012; Bradford et al., 2015). Contrary to full-length Pan1 and Sla2, we detected FRET of Pan1(1-1050) with Clc1 and Chc1 (1.4% and 1.2%; *p*=0.004 and *p*=0.001 against Pan1-Clc1 and Pan1-Chc1 pairs, respectively; *n*=12 cells), and between Sla2dTHATCH and Clc1 (1.4%; *p*=0.009 against Sla2-Clc1; *n*=14 cells) (Figure 3B). These data strongly indicate that the C-termini of Pan1 and Sla2, and indirectly their proximity partner the Sla1 N-terminus, are located at the cytoplasmic site of the clathrin lattice, being probably more than 10 nm offset.

#### Key coat proteins transverse the clathrin lattice in the membrane-cytoplasm direction

As protein shortening showed to be a powerful tool for mapping the coat organization, we used it further to increase the precision of our map. First, we focused on the absence of FRET between the Sla2 and Pan1 C-termini, which were both identified as members of the cytoplasmic proximity network. We measured FRET between the Sla2 C-terminus and Pan1(1-1050) and detected very high FRET value (8.5 ± 0.9%). We also constructed a less shortened Pan1 C-terminal truncation Pan1(1-1303) and again detected FRET with the Sla2 C-terminus. Its lower value (4.8 ± 0.5%; *p*<0.001, *n*=15 cells) suggested that the Pan1 C-terminus is the most remote part of the cytoplasmic proximity network (Figure 3B).

Finally, we tested the proposed Pan1 orientation by using a functional Pan1(402-1480) truncation, which N-terminally shortened Pan1 after its first EH domain (Bradford et al., 2015). We compared FRET values of this N-terminal truncation and the Pan1 N-terminus towards both clathrin subunits. Although FRET of Pan1(402-1480) and the Pan1 N-terminus with Chc1 were similar, only Pan1(402-1480) showed FRET with Clc1 (1.8%; *p*=0.035, *n*=11 cells), suggesting that this region is closer to the cytoplasmic site of the clathrin lattice than the complete Pan1 N-terminus (Figure 3B).

To sum up, the detected proximities of Pan1 and Sla2 full-length and truncated proteins strongly support their following topology: both proteins have their N-termini between the plasma membrane and the clathrin lattice, their mid parts are spanning through the lattice, and their C-termini protrude deeply into the cytoplasm, especially in case of Pan1. The same layout can be inferred for Sla1 protein, but with the opposite orientation of its termini (Figure 3A and B).

#### FRET proximities of Gts1 suggest its orientation towards the plasma membrane

The comparison of FRET values of the N-and C-termini of endocytic proteins (e.g. End3) greatly helped to orient them in the coat map (see above). They were made on the assumption that if the N-and C-terminus of a protein show significantly different FRET to the same proximity partner, these differences most probably mirror their relative distances to this partner. We applied this approach to further map the ArfGAP protein Gts1 inside the membrane-proximal protein network. The presence of the Gts1 C-terminus in this network was firmly established by its specific FRET with all network members. FRET of the Gts1 N-terminus with the Sla1 C-terminus and, distinctly to all other proteins, with the membrane-associated Sla2 N-terminus indicated that the Gts1 N-terminus is also part of this network, being potentially more adjacent to the membrane (Figure 3A). Nevertheless, to locate the Gts1 N-terminus more precisely, we needed to test its proximity to the N-termini of Pan1 and Sla2, for which we needed a second functional N-terminal tag. We therefore introduced new FRET fluorophore pairs mTurquoise2-mNeonGreen and mNeonGreen-mScarlet-I, which can be both used for functional N-terminal tagging (data not shown). We then measured FRET of both Gts1 termini tagged with mScarlet-I to N-terminus of Pan1 or Sla2 tagged with mNeonGreen (Figure 3C). While both Gts1 termini showed similar FRET towards the Pan1 N-terminus (3.2% and 3.7%), FRET with the Sla2 N-terminus was much higher for the Gts1 N-terminus than for the Gts1 C-terminus (6.1% vs. 1.4%; *p*=0.002, *n*=13 cells). These data together with FRET values of Gts1 ends with the Sla1 C-terminus (1.7% and 7.7% for the Gts1 N-and C-terminus, respectively; *p*<0.001, *n*=14 cells) strongly suggest that the Gts1 N-terminus is placed closer to the plasma membrane than the Gts1 C-terminus. Indirectly, these data also imply a more membrane-proximal location of the Pan1 N-terminus than the Sla1 C-terminus, though probably still more remote than the Gts1 and Sla2 N-termini (Figure 3A and C). In agreement, strong FRET was detected between N-termini of Pan1 and Sla2 tagged with mNeonGreen and mScarlet-I and *vice versa* (4.6% and 5.3%, respectively).

### FRET proximity screen detects cytoplasmic assemblies of endocytic coat proteins

Fluorescence cross-correlation spectroscopy (FCCS), another powerful microscopy technique for detection of molecular interactions *in vivo*, was recently applied to search for assemblies of endocytic proteins in the yeast cytoplasm (Boeke et al., 2014). We tested if our proximity screen could also identify cytoplasmic subcomplexes of endocytic coat-associated proteins similarly to FCCS. Indeed, we detected all three cytoplasmic interactions between Syp1-Ede1, Pan1-End3, and Las17-Sla1 found by FCCS (Boeke et al., 2014) by their cytoplasmic FRET (Table 1). Strikingly, while FRET between Syp1-Ede1 and Pan1-End3 pairs persisted at endocytic sites, FRET between Las17 and Sla1 was specifically detected only in the cytoplasm. This was the case for C-terminally tagged Las17 and Sla1, but also for Las17 and Sla1 proteins tagged with mTurquoise2 and mNeonGreen either on the N-and C-terminus, respectively, or on their N-termini (Table 1), which are supposedly closer to the proposed Sla1-Las17 interaction surface (Feliciano and Di Pietro, 2012). These data indicate a substantial rearrangement of the Sla1-Las17 complex at the endocytic site happening already before the onset of actin polymerization.

**Table 1.** Cytoplasmic assemblies of coat-associated proteins detected by FRET

| Protein pair<sup>a</sup> |  | FRET%<sup>b</sup> |  |
| --- | --- | --- | --- |
|  |  | cytoplasm | endocytic sites |
| Syp1• | Ede1• | 2.7 ± 0.6 (38) | 3.3 ± 0.6 (38) |
| •End3 | Pan1• | 4.2 ± 1.8 (20) | 5.8 ± 1.0 (20) |
| Las17• | Sla1• | 1.5 ± 2.2 (13) | 0.1 ± 0.7 (13) |
| •Las17 | Sla1• | 8.5 ± 1.3 (30) | 0.0 ± 2.3 (31) |
| •Las17 | •Sla1 | 9.8 ± 2.9 (24) | 0.7 ± 2.5 (36) |

<sup>a</sup> Syp1-GFP+Ede1-mCherry, GFP-End3+Pan1-mCherry, Las17-GFP+Sla1-mCherry, mTurquoise2-Las17+Sla1-mNeonGreen, and mTurquoise2-Las17+mNeonGreen-Sla1 strains were used for the analysis. • denotes the position of the fluorophore tag on the protein.
<sup>b</sup> Mean FRET percentages with 95% confidence intervals and number of analyzed cells (in parentheses) are shown.

### Rearrangements of coat proteins during membrane invagination

The reorganization of certain parts of the endocytic coat is predicted during the membrane invagination steps of endocytosis (Picco et al., 2015; Sochacki et al., 2017; Mund et al., 2018). Such changes cannot be traced by our screen, which specifically targets endocytic coats on the still flat plasma membrane. To probe if coat proteins get rearranged during membrane invagination, we measured FRET of selected proteins pairs at endocytic sites sorted according to their endocytic stage. For that we additionally tagged protein Abp1 as a marker of actin-dependent invagination steps of endocytosis (Kukulski et al. 2012), and worked on fixed cells as recently established for super-resolution microscopy of yeast endocytosis (Mund et al., 2018). We focused on three conceivable protein-protein rearrangements. We followed detected FRET proximities between Ent1-Sla1 and Pan1(1-1303)-Sla2 C-termini, which could be eventually lost during membrane invagination; and looked for a potential presence of FRET proximity between the Ent1-Sla2 C-termini around the same time. During invagination the Ent1 and Sla2 C-termini could colocalize at the cytoplasmic area of the coat by their functional binding to the actin cytoskeleton (Skruzny et al., 2012), while the Sla1 and Pan1(1-1303) C-termini could separate to an outer rim of the coat as suggested by recent studies (Picco et al., 2015; Sochacki et al., 2017; Mund et al., 2018) (see scheme in Figure 4A). We performed acceptor photobleaching in strains expressing the indicated protein pairs tagged with mTurquoise2 and mNeonGreen together with Abp1-mScarlet-I, and analyzed FRET in individual endocytic sites sorted according to the presence of invagination marker Abp1. In agreement with our screen, we observed FRET between Ent1-Sla1 and Pan1(1-1303)-Sla2 at the early endocytic sites absent of Abp1. Similar FRET was observed for the Pan1(1-1303)-Sla2 pair at endocytic sites decorated with Abp1, suggesting that the Pan1(1-1303) region and the Sla2 C-terminus remain proximal also during membrane invagination (Supplementary Figure 2). Notably, FRET between the Ent1-Sla1 C-termini was slightly, but significantly, decreased in the endocytic sites containing Abp1 (Figure 4B). This is indicative of partial separation of the Ent1 and Sla1 C-termini during invagination. Most strikingly, though the Ent1 and Sla2 C-termini showed no FRET in our screen or at the endocytic sites absent of Abp1, a high FRET between these protein termini was detected at invaginating endocytic sites containing Abp1 (Figure 4C). This suggests that initially separated Ent1 and Sla2 C-termini join together during actin-dependent membrane invagination.

**Figure 4.**
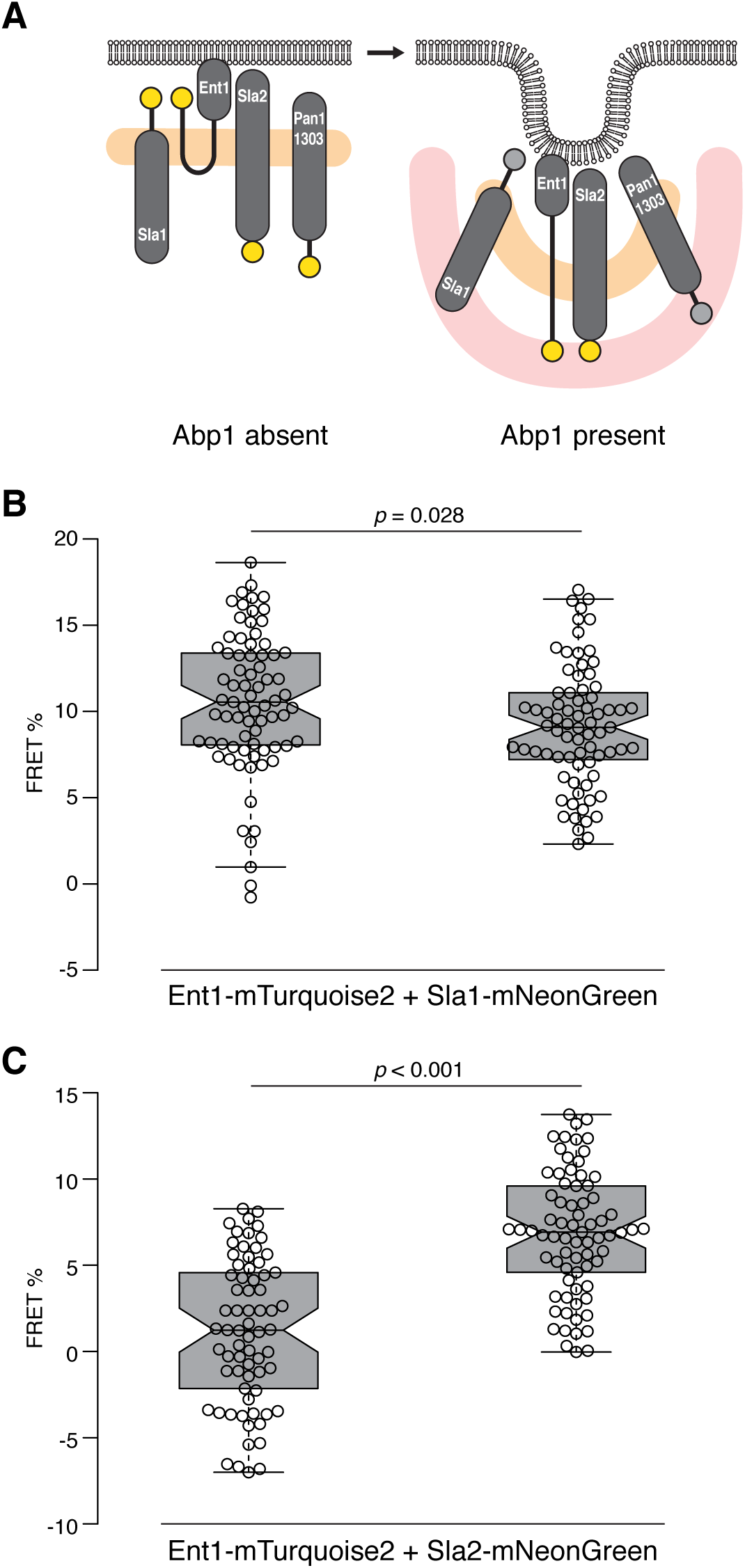
Rearrangements of endocytic coat proteins during membrane invagination. (A) Scheme of suggested topology of Sla1, Ent1, Sla2 and Pan1(1-1303) proteins on the flat (left) and invaginated (right) plasma membrane. Natively unfolded parts of the proteins are shown as thick black lines. The C-termini of proteins showing FRET are depicted as yellow circles; the C-termini separated from their FRET partners during invagination are shown as gray circles. The position of the clathrin lattice and polymerizing actin cytoskeleton are shown as a pale orange and red zone, respectively. **(B-C)** Partial loss of FRET between Ent1-mTurq-uoise2 and Sla1-mNeonGreen **(B)** and occurence of FRET between Ent1-mTurquoise2 and Sla2-mNeonGreen **(C)** during endocytic membrane invagination. FRET values (in %) of indi-vidual endocytic patches (n=74, 75; 69, 73 for **(B)** and **(C)**, respectively) sorted according the absence/presence of Abp1 are shown as box plots. Center, top and bottom lines of box plots show the medians, the 25th and 75th percentiles of individual datasets, respectively. Whiskers extend 1.5 times the interquartile range from the 25th and 75th percentiles. Notches indicate 95% confidence intervals of the medians. See also Supplementary Figure 2.

### Binding dynamics of coat-associated proteins at the endocytic site

Having the spatial organization of the endocytic coat resolved, we also wanted to analyze binding dynamics of its protein components. Specifically, we used fluorescence recovery after photobleaching (FRAP) to follow if coat-associated proteins are stably or dynamically bound at the endocytic site thus showing slow or fast protein exchange, respectively, of their in-site bleached molecules. We followed FRAP of Ede1, Apl1, Yap1801, Gts1, Bzz1, Lsb3, Las17, and Vrp1 tagged with mNeonGreen similarly as it was previously done for other coat-associated proteins tagged with EGFP (Skruzny et al., 2012). As shown in Figure 5 and Movie 2, Apl1, Yap1801, and Gts1 proteins showed very low fluorescence recovery, similar as it was found for Ent1/2, Sla2, Pan1, Sla1 and End3 (Skruzny et al., 2012). Much higher recovery was detected for Ede1, Las17, Lsb3 and Vrp1 proteins, though the recovery of low-abundant Vrp1 seemed to be lower. Finally, a very high recovery was observed for the F-BAR protein Bzz1. In summary, FRAP experiments showed that while endocytic adaptors Apl1 and Yap1801, and Gts1 protein are stably bound at the endocytic coat, the endocytic priming proteins Ede1 and Syp1, and actin regulators Las17, Lsb3, and Bzz1 associate with the endocytic machinery in a dynamic manner.

**Figure 5.**
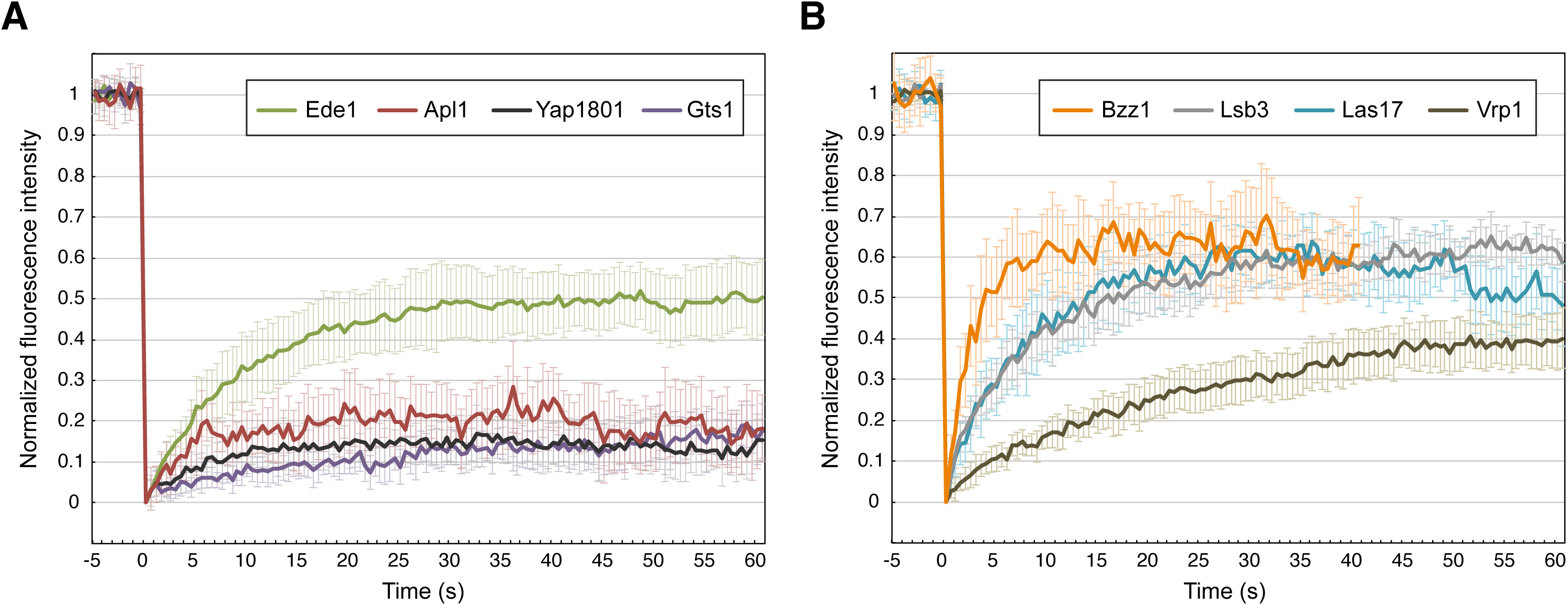
FRAP analysis of indicated coat-associated proteins at the endocytic site. Ede1, Apl1, Yap1801, Gts1, Bzz1, Lsb3, Las17, and Vrp1 proteins tagged with mNeonGreen were photobleached in individual endocytic sites of LatA-treated cells and fluorescence recovery was followed every 0.5 s for 60 s (40 s in case of the highly dynamic Bzz1 protein). The curves represent means ± 95% confidence intervals (n=9–17).

## Discussion

To understand the mechanism and regulation of endocytic vesicle formation, the knowledge of its underlying protein machinery is of utmost importance. Here, we describe the molecular organization of the endocytic coat, the main structural and functional element of the endocytic site, using a robust and reliable FRET technique. We found that coat proteins are organized in distinct functional layers situated on both sides of the clathrin lattice, and that key endocytic factors interconnect these layers in the membrane-cytoplasm direction. Moreover, we describe rearrangements of the endocytic coat during membrane invagination and characterize binding dynamics of involved proteins.

We focused on the endocytic coat of budding yeast *Saccharomyces cerevisiae*, the proteins of which are all, or almost all, known. In comparison to mammalian endocytic coats, which contain many lately evolved cargo-, tissue-and splicing-specific proteins, the yeast endocytic coat consists of only 22 proteins covering many important families of endocytic adaptors, scaffolds and coat-associated actin regulators (Weinberg and Drubin, 2012; Goode et al., 2015; Lu et al., 2016). Notably, the majority of proteins are represented by a single paralog with the exception of three duplicates: CALM/AP180 proteins Yap1801/2, epsins Ent1/2 and SH3YL1 homologs Lsb3/4. Most importantly, our 18 studied proteins are shared by many fungal and animal species and potentially constitute the evolutionary most conserved factors of the endocytic machinery (Dergai et al., 2016). The yeast endocytic coat could therefore represent a common functional core of a general endocytic coat, and its uncovered protein architecture is likely to be shared by many organisms. Future studies of endocytic coats from other species will hopefully support this notion.

The proposed organization of the yeast endocytic coat on the flat plasma membrane is depicted in Figure 6. Interacting endocytic pioneering proteins Ede1 and Syp1 (Reider et al., 2009) were found proximal to each other, but not to other proteins. This can be a consequence of our focus on the late endocytic coat, where Ede1/Syp1 interactions can already be replaced by a stable protein network provided by similar domains of Pan1, End3, and Sla2 proteins. In agreement, our FRAP data show a similar dynamic association of Ede1 and Syp1 with the endocytic site, in contrast to a stable binding of Pan1, End3, and Sla2 (Figure 5; Skruzny et al., 2012). Ede1 and Syp1 proteins are thus likely to dynamically localize outside of the coat as suggested by their recent super-resolution imaging (Mund et al., 2018) and their incomplete colocalization with other proteins in our screen (Supplementary Figure 1).

**Figure 6.**
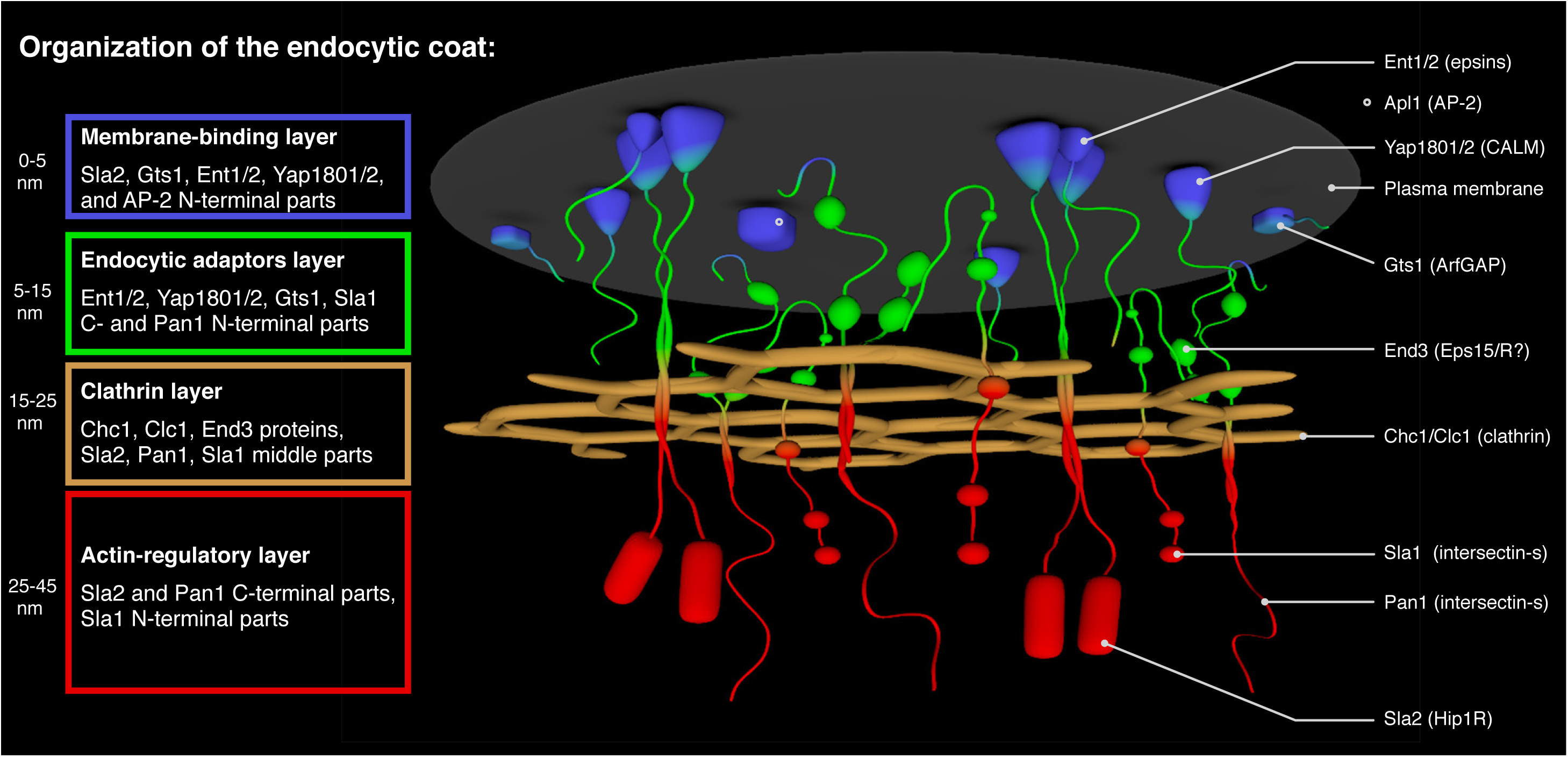
Model of the protein architecture of the endocytic coat. Individual coat layers together with their constituents and their approximate distances from the plasma membrane are listed on the left. Legend to individual depicted proteins (human homo-logs in parentheses) is shown on the right. Regulators of actin polymerization Las17, Vrp1 and Bzz1, and endocytic pioneering proteins Ede1 and Syp1, which were recently shown to localize in membrane-proximal rings outside of depicted endocytic coat, are not shown for clarity. See the text for further details.

As suggested by their extensive FRET proximities and similar low FRAP recoveries (Figures 3 and 5; Skruzny et al., 2012), a stable structural core of the yeast endocytic coat is made of adaptors AP-2 (represented by Apl1 subunit), Yap1801/2, Ent1/2 and Sla2; endocytic scaffolds Pan1, End3, Sla1 and ArfGAP protein Gts1. Proximity mapping with clathrin subunits as topology markers then sorted these factors into two distinct spatial layers located above and below the clathrin lattice. As expected, prototypical adaptor proteins AP-2, Yap1801/2 and Ent1/2 localize between the plasma membrane and clathrin, so we named this layer accordingly, the endocytic adaptors layer. They share this region with Gts1, the N-termini of Sla2, Pan1 and End3, and the Sla1 C-terminus. FRET measurements with the truncated Sla2 and Pan1 proteins then established the second layer, which surprisingly localized on the cytoplasmic site of the clathrin lattice. This layer consists of the C-termini of Sla2, Pan1, and End3; and the Sla1 N-terminus. Sla2, Pan1 and Sla1 proteins therefore span through the clathrin lattice and connect both layers above and below it. As the C-termini of Sla2 and Pan1, and the Sla1 N-terminus are extensively involved in either binding (Sla2, Pan1) or regulation (Pan1, Sla1) of the actin cytoskeleton at the endocytic site, we named this area, the actin regulatory layer. Based on FRET results of shortened Sla2 and Pan1 proteins, we propose that the actin regulatory layer (especially the Pan1 C-terminus) extends more than 10-20 nm from the clathrin lattice. If we assume that these distances are also kept in the membrane plane of the coat and during membrane invagination, the respective Sla2, Pan1 and Sla1 parts are ideally placed not only for their binding and regulation of the actin cytoskeleton, but also for the proposed communication of Pan1 and Sla1 with actin nucleators Myo3/5 and Las17, respectively, which are localized at the outer, membrane-proximal ring around the coat (Barker et al. 2007; Picco et al., 2015; Mund et al., 2018). Although limited to FRET data of Sla2 and Gts1 N-termini, we can also discern the membrane-proximal part of the adaptors layer as the membrane-binding layer. This layer consists of the N-termini of Sla2 and Gts1, and supposedly all N-terminal membrane-binding domains of adaptors AP-2, Yap1801/2 and Ent1/2; and can be characterized by the absence of FRET to clathrin. The presence of the N-terminal ArfGAP domain and the central ALPS motif, involved in membrane biochemistry of Gts1 (Smaczynska-de Rooij et al., 2008), are supportive for the placement of the Gts1 N-terminus in this layer (Figure 3C).

Similarly to Ede1 and Syp1, we did not detect any adjacent proteins to the actin regulators Las17, Vrp1 and Bzz1, apart from the expected proximities between them (Naqvi et al., 1998; Soulard et al., 2002). Based on recent evidence (Mund et al., 2018), we expect these proteins to localize in the outer ring around the endocytic coat, binding there, according to our FRAP studies, in a dynamic manner.

On top of a detailed description of the endocytic coat architecture, our FRET experiments also identified two significant coat rearrangements occurring during membrane invagination. Firstly, we observed a decrease of FRET between the Sla1 and Ent1 C-termini indicative of their partial separation. This is in line with recently suggested relocalization of Sla1 and its human homolog intersectin-s to the outer rim of the coat during invagination (Picco et al., 2015; Sochacki et al., 2017). However, a small drop in FRET suggests that this Sla1-Ent1 segregation is either not large or occurs only within a sub-pool of Sla1-Ent1 molecules. Both notions are consistent with a recent super-resolution study showing i) only small changes (around 5 nm) of the average position of Sla1 functional partner Pan1 during endocytosis, and ii) only a small fraction of endocytic sites with rim-organized Sla1 (Mund et al., 2018). Secondly, we observed presence of FRET proximity between the Ent1 and Sla2 C-termini specifically occurring during membrane invagination. The Ent1 C-terminus, which contains both clathrin-and actin-binding motifs (Wendland et al., 1999; Skruzny et al., 2012), thus seems to relocate from the adaptors layer of the coat to its actin layer after the onset of actin-driven membrane invagination. This supports the proposed role of the Ent1 C-terminus in harnessing the force of actin polymerization to productive membrane invagination (Skruzny et al., 2012; Messa et al., 2014). Future studies will hopefully provide more cases to these pivotal examples of functional protein rewiring during endocytosis.

We also analyzed the acquired coat map with respect to known functional, binding and regulatory motifs of the involved proteins (Supplementary Figure 3A). As formation of the endocytic coat requires interaction of membrane-bound adaptors with clathrin, cargo and each other, the exclusive presence of all known clathrin-, cargo-and adaptor-binding motifs and domains in the adaptors and clathrin layers is perhaps not surprising. More intriguingly, all predicted phosphorylation sites of coat-associated kinases Ark1/Prk1 (AAK1/GAK1 in human) are also located in the adaptors layer, suggesting that this layer is the primary place of action for these uncoating kinases. Notably, not only the actin-binding and actin-nucleating domains of Sla2 and Pan1, respectively, but also the Pan1/Sla1 polyprolin motifs (PP) and the Sla1 SH3 domains colocalize in the actin-regulatory layer of the coat. This is consistent with the proposed formation of a multivalent SH3-PP network made by Pan1, Sla1 and actin regulators Las17, Vrp1, Myo3/5 and Bbc1, necessary for an efficient regulation of actin polymerization at the endocytic site (Sun et al., 2017; Mund et al., 2018). Moreover, the cytoplasmic location of the SH3-PP network is favorable for its potential role in the phase separation, recently suggested to aid in endocytic membrane invagination (Bergeron-Sandoval et al., 2017).

Finally, we compared the coat proximity map with known interactome of its protein constituents. We found that 60% (27 of 45) of previously described interactions were recovered by our proximity screen. As shown in Supplementary Figure 3B, the missed interactions involved mostly interactions of Ede1, Syp1, Las17, and Lsb3 proteins, which might not be detected for reasons already discussed. Most importantly, our screen generated a large and robust dataset of 19 protein pair proximities, which were not mirrored by respective protein-protein interactions. Some of them may therefore represent yet-to-be characterized functional protein-protein interactions or subcomplexes of the endocytic coat.

Altogether, our study provides a highly-resolved spatial, regulatory and functional map of the conserved endocytic coat. We believe that its comprehensive description will foster mechanistic understanding of the endocytic process and serve as a framework for characterization of other multiprotein assemblies involved in endocytosis and other membrane reshaping/trafficking cellular pathways.

## Supporting information

Skruzny et al-Movie 1

Skruzny et al-Movie 1

## Acknowledgements

We thank A. Zori Comba for 3D model in Figure 6, and M. Abella and S. M. Murray for critical reading of the manuscript. This work was funded by Deutsche Forschunsgemeinschaft (DFG) Research Grant SK 305/1-1.

## Author contributions

Conceptualization, Project Administration, M.S.; Methodology and Resources, M.S., G.M.; Investigation and Formal Analysis, M.S., E.P., S.G.; Writing - Original Draft, M.S.; Writing - Review & Editing, M.S. and V.S.; Supervision and Funding Acquisition, M.S. and V.S.

## Experimental Procedures

### Yeast strains and media

Standard yeast media and protocols were used to manipulate yeast strains listed in Supplementary Table 3. The N-and C-terminal tagging/truncation of yeast proteins was made by homologous recombination of respective genes with PCR cassettes previously described (Janke et al., 2004, Khmelinskii et al., 2011) or constructed in the lab. For microscopy, strains were grown to a log phase in a low fluorescence SD-Trp medium (prepared from LoFlo YNB, Formedium). Cells were attached to Concanavalin A-coated (0.1 μg/ml, Sigma) 8-well glass coverslips (ibidi) and observed at 25 °C. Where indicated, cells were incubated with 200 μM Latrunculin A (Enzo Life Sciences) for 10 min followed by 10-50 min of imaging. Cell fixation with 16% formaldehyde (Thermo) was performed on cells attached to 8-well coverslips as described previously (Mund et al., 2018).

### FRET microscopy

Acceptor photobleaching was performed using a wide-field Eclipse Ti-E inverted fluorescence microscope (Nikon) equipped with X-Cite Exacte LED light source, Perfect Focus System (PFS) and NIS-Elements AR software (4.40; Nikon). Images were acquired with Nikon 60x Plan Apo NA 1.45 oil immersion objective with 1.5x tube lens and iXon 897-X3 EM-CCD camera (Andor) with EM gain set up to 250. Following filters were used to acquire mTurquoise2 (Ex 436/20, Em 480/40), GFP (Ex 470/40, Em 525/50), mNeonGreen (Ex 504/12, Em 542/27) and mCherry/mScarlet-I (Ex 585/29, Em 647/57) fluorescence (Chroma, Semrock). Two acceptor-and three to five donor-channel images were taken before photobleaching of the acceptor by 3-5 s pulse of 150 mW 593 nm (mCherry/mScarlet-I) or 515 nm (mNeonGreen) solid-state laser (Acal BFi) followed by three to five donor-and two acceptor-channel images. In general, the more abundant protein of a protein pair was used as the FRET acceptor. Several protein pairs were also analyzed with swapped fluorophore tags giving qualitatively same FRET results as the originally tagged protein pair (Supplementary Figure 1 and data not shown). Images were analyzed with ImageJ software (Schneider et al., 2012). Images were subtracted of general background and endocytic patches of each photobleached cell were then manually selected by polygon selection tool. FRET efficiency (calculated as percentage increase in donor fluorescence after acceptor photobleaching) was calculated using FRETCalc plugin (Stepensky, 2007) with the intensity threshold set up to the level of cytoplasmic fluorescence of the analyzed cell. At least 6 cells (each contributing by multiple endocytic patches) of two independent clones were used to calculate mean FRET efficiency and 95% confidence intervals of respective protein pair. FRET efficiency values were finally corrected by subtraction with FRET values of identically acquired and processed respective donor-only strains (showing zero or slightly negative FRET due to a mild structural photobleaching of donor during acquisition). Welch’s form of t-test was used to calculate statistical differences between individual FRET datasets.

### FRAP microscopy

FRAP microscopy was performed using Visitron wide-field fluorescence microscope (Visitron Systems) based on Nikon Eclipse Ti-E inverted system equipped with PFS, CoolLED pE-4000 light source, and VisiFRAP System operated by VisiView software (3.3.0.6). Images were acquired with Nikon 100x Apo TIRF NA 1.49 oil immersion objective and iXon Ultra-888 EM-CCD camera (Andor). Chroma TIRF ET 514 nm filter set was used to acquire mNeonGreen fluorescence. Photobleaching was achieved by 20-100 ms pulse of 100 mW 515 nm laser (used at 1-5% power) and fluorescence recovery was followed for 1-2 min with 500 ms frame rate. FRAP was analyzed as described previously (Skruzny et al., 2012).

## Supplementary Movies Legends

**Movie 1. FRET between indicated protein pairs observed as an increase of GFP donor fluorescence after mCherry acceptor photobleaching.** Fluorescence of GFP FRET donor attached to the indicated protein was observed before and after photobleaching of mCherry FRET acceptor. Five frames of 500-1000 ms exposure acquired before and after photobleaching are shown.

**Movie 2. FRAP of mNeonGreen-tagged Ede1, Apl1, Yap1801, Gts1, Bzz1, Lsb3, Las17, and Vrp1 proteins in LatA-treated wild-type cells.** Indicated proteins were photobleached at the endocytic sites marked with arrows (at time 0) and their fluorescence recovery was followed for 60 s with 500 ms frame rate.

**Supplementary Figure 1.**
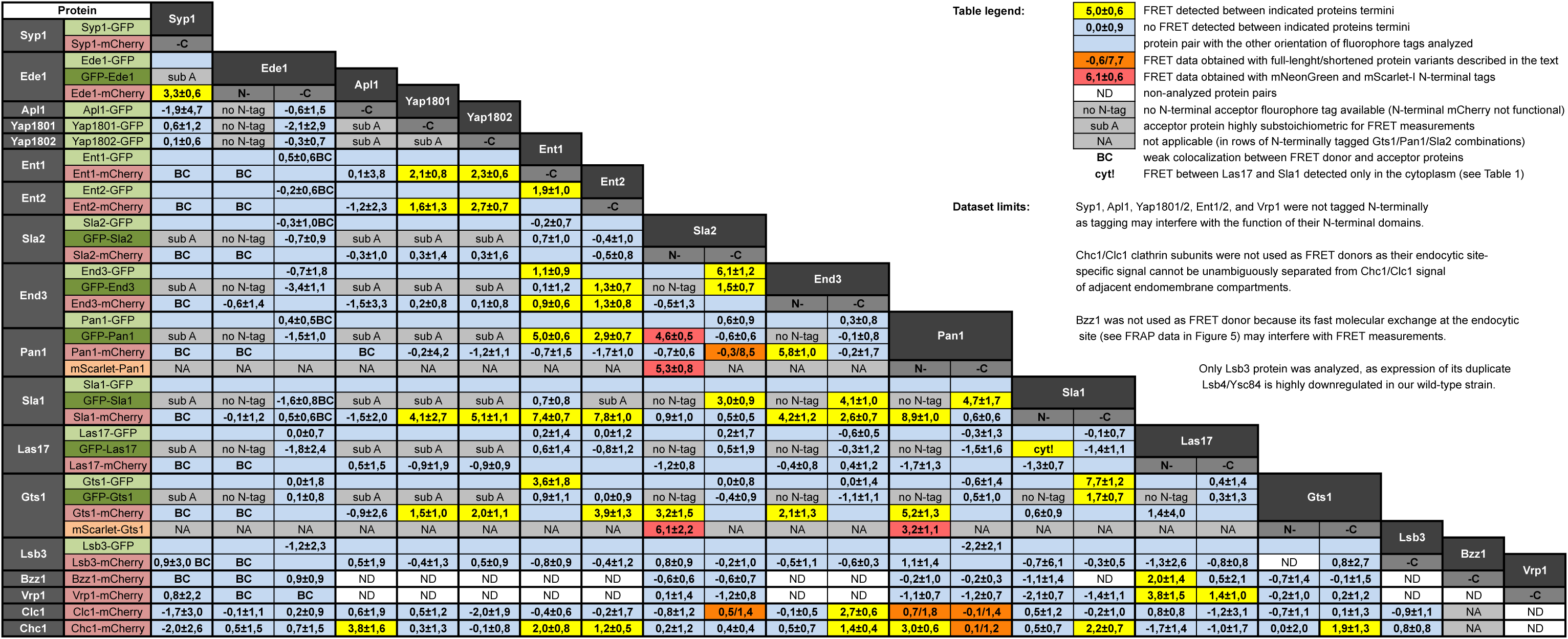
The complete overview of FRET-based protein proximity screen of the yeast endocytic coat. Mean FRET efficiencies (in % with 95% confindence intervals) calculated from multiple endocytic sites of at least 6 cells are shown for individual protein pairs. See the text on the top right for detailed description of the dataset.

**Supplementary Figure 2.**
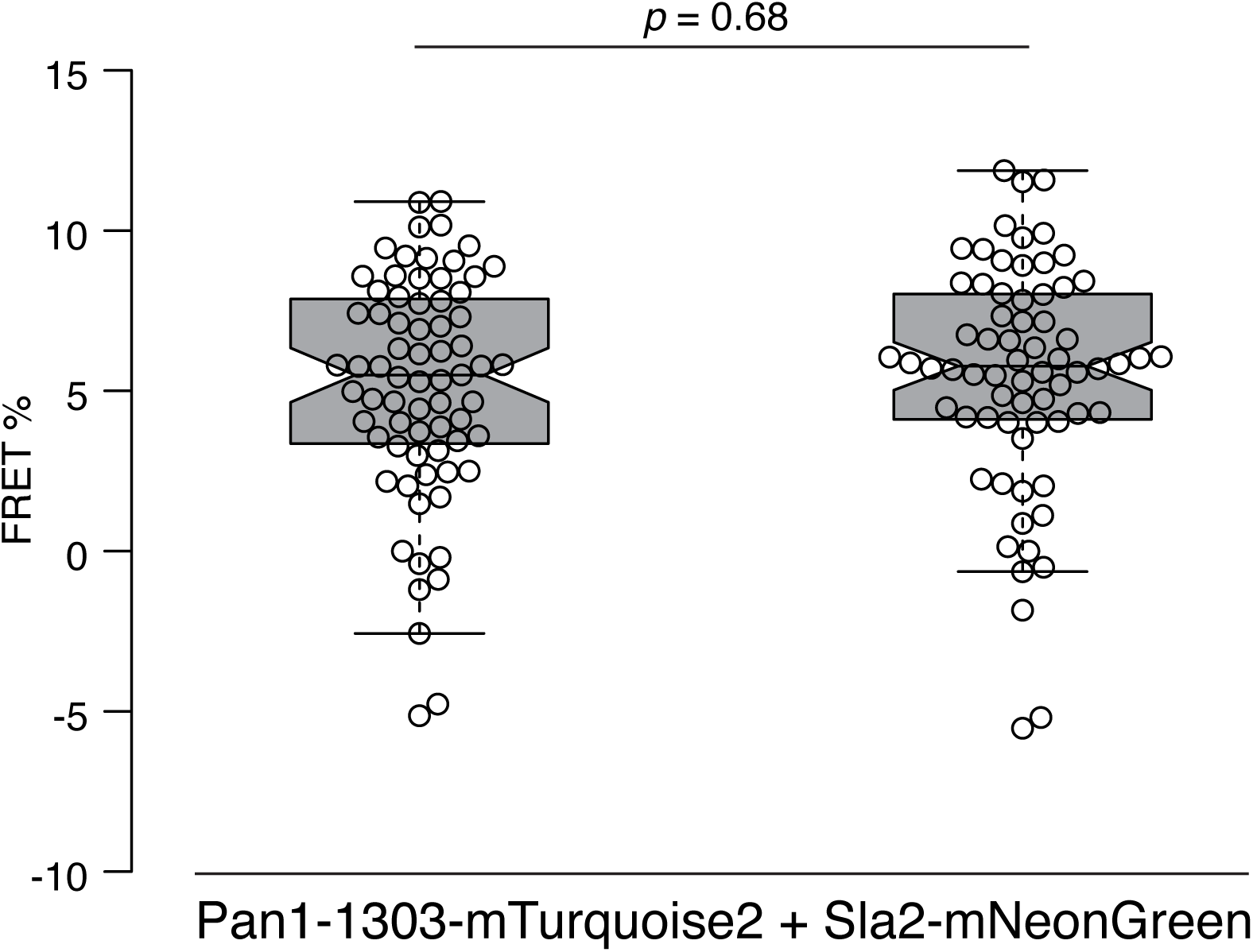
FRET proximity between Pan1(1-1303)-mTurquoise2 and Sla2-mNeonGreen does not change during membrane invagination. FRET values (in %) of 71 or 68 endocytic patches containing or absent of Abp1, respectively, are shown as box plots. Center, top and bottom lines of box plots show the medians, the 25th and 75th percentiles of individual datasets, respectively. Whiskers extend 1.5 times the inter-quartile range from the 25th and 75th percentiles. Notches indicate 95% confidence intervals of the medians.

**Supplementary Figure 3.**
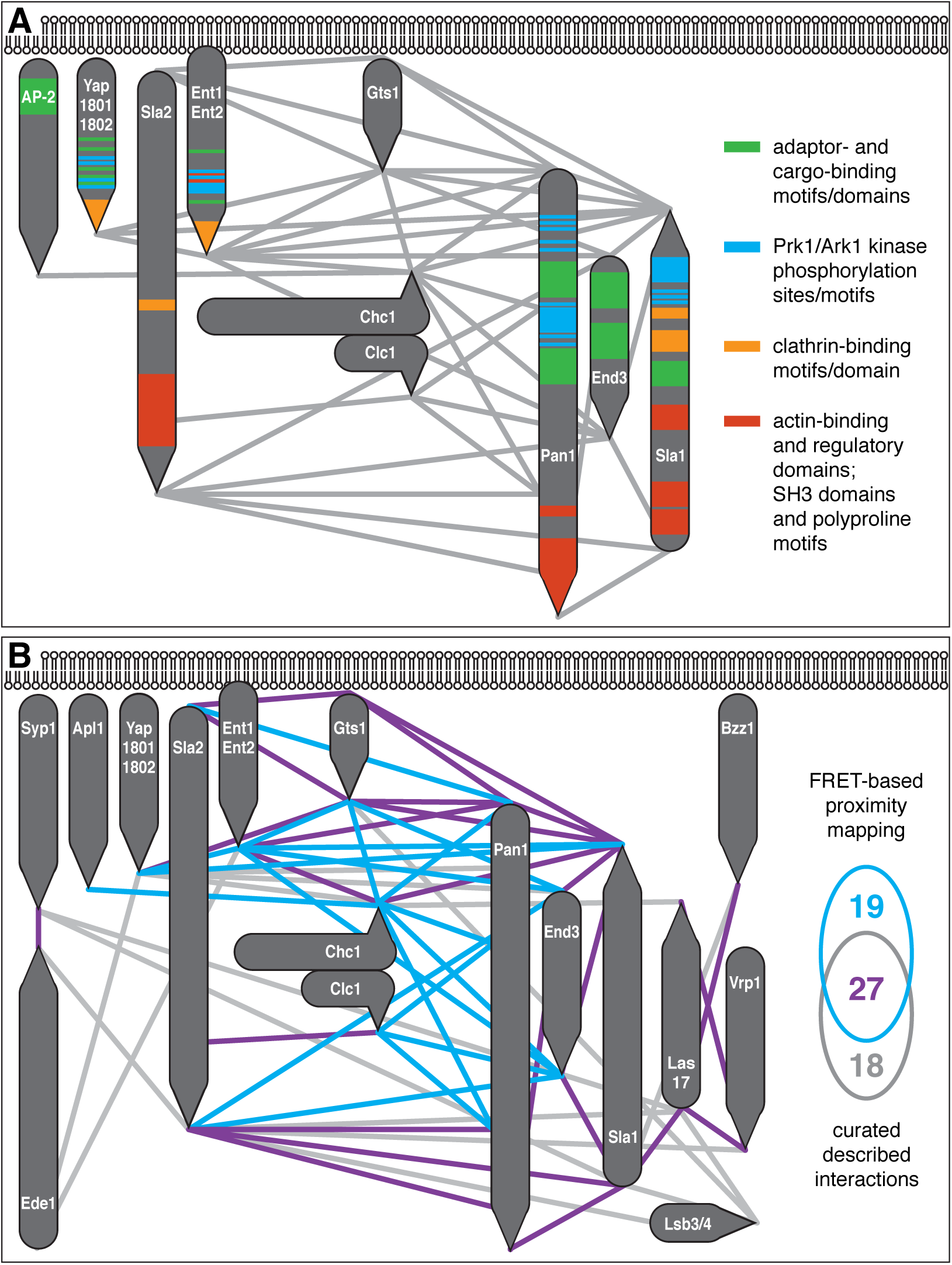
Analysis of functional, binding and regulatory moieties of the endocytic coat architecture. Topology of functional, binding and regulatory motifs and domains in the proposed coat architecture. Known motifs and domains of indicated coat proteins were classified according to their main function and color-coded as shown on the right. Ark1/Prk1 phosphorylation sites were taken from (Huang et al., 2003). **(B)** Comparison of the FRET proximity mapping with previously described interactions of studied endocytic proteins. Physical interactions of indicated proteins (observed in more than a single study) were taken from Saccharomyces Genome Database as per October 2018. FRET proximities mirroring previously detected protein interactions are shown as violet lines. Proximities not found as interactions and *vice versa* are shown as blue and gray lines, respectively.

